# Comparative genomics of five *Valsa* species gives insights on their pathogenicity evolution

**DOI:** 10.1101/2022.05.17.492390

**Authors:** Guangchao Sun, Shichang Xie, Lin Tang, Chao Zhao, Mian Zhang, Lili Huang

## Abstract

*Valsa* is a genus of ascomycetes fungi within the family Valsaceae that includes many wood destructive pathogens. The species such as *Valsa mali* and *Valsa pyri* that colonize fruit trees are threatening the global fruit production. Rapid host adaptation and fungicide resistance emergence are the main characteristics that make them devastating and hard to control. Efficient disease management can be achieved from early infection diagnosis and fungicide application, but lack of understandings of their genetic diversity and genomic features that underpin their pathogenicity evolution and drug resistance is essentially impeding the progress of effective and sustainable disease control. Here, we report genome assemblies of *Valsa malicola, Valsa persoonii* and *Valsa sordida* which represents close relatives of the two well known *Valsa mali* and *Valsa pyri* that cause canker disease with different host preferences. Comparative genomics analysis revealed that segmental rearrangements, inversions and translocations frequently occurred among *Valsa* spp. genomes. Genes identified in highly active regions exhibited high sequence differentiation and are enriched in membrane transporter proteins involved in anti-drug and nutrient transportation activities. Consistently, we also found membrane transporter gene families have been undergoing significant expansions in *Valsa* clade. Furthermore, unique genes that possessed or retained by each of the five *Valsa* species are more likely part of the secondary metabolic (SM) gene clusters which suggests SM one of the critical components that diverge along with the evolution of <I>Valsa</I> species. Repeat sequence content contributes significantly to genome size variation across the five species. The wide spread AT-rich regions resulted from repeat induced point C to T mutation (RIP) exhibited a specific proximity to secondary metabolic gene clusters and this positional proximity is correlated with the diversification of SM clusters suggesting a potential companion evolution between repeat sequence and secondary metabolism cluster. Lastly, we show that *LaeA*, the global regulator of secondary metabolic gene cluster, exhibiting diverged manner of regulation on the expression of clusters in vegetative and invasive mycelia of the devastating *V. mali* indicating the complexity of secondary metabolism in fungal species.

## Introduction

*Valsa* spp., a fungal genus of Ascomycota phylum, includes several destructive woody canker pathogens that infect apple, pear, poplar, cherry peach and many other economically or ecologically important rosids [1]. For example, *Valsa mali*, a causal agent for apple canker disease, has become notorious for causing huge yield losses of seasonal apple production in eastern Asia [2, 3]. In areas with severe incidence, apple gardens can be deteriorated and it takes years to recover. Recent efforts taken to understand the molecular mechanisms underlying the pathogenecity have paved a way towards effective strategies for disease control [4, 5]. However, likely due to a highly deversed genetic pool, new pathovars that are capable of circumventing current disease control strategies frequently emerge. Several recent comparative genomics studies on *Valsa mali* and *Valsa pyri* suggested genomic adaptation substantially contributed to their virulence [6, 7]. A significant change in the content of pathogenicity related genes involved in cell wall degradation, host immunity suppression, secondary metabolite biogenesis and repeat sequences were observed since their divergence from the common ancestor 4 million years ago [6]. Moreover, expansion of secondary metabolism gene clusters via gene gain and transposon insertions in *Valsa mali* genome is prevalent in comparison to *Valsa pyri*. It has been hypothesized that as genetic consortia, metabolic gene clusters may encode a variety of regulatory mechanisms to keep the persistence and protect their host fungi while contribute to ecological adaptation of fungi [1]. For example, a subset of cutinase from *M. orzyae* and its relatives was phylogenically close to actylxylase which is responsible for xylan degradation instead of their original substrate cutin. This might imply a selection on these gene sets for woody host adaptation. In *Valsa spp*., a similar pattern of selection on pectinase gene family was shown to be important for infection and host adaptation [6, 8].

Valsa species exhibited obvious host preference as observed in previous and current study (Figure 1A). As mentioned hereinbefore, *V. pyri* diverged from the common ancestor with *V. mali* at 4 million years ago while currently showing preferential infection on pears even though it still colonizes apple trees (Figure 1A). This fast host switch might be a consequence of population competition [9] and cannot be achieved without active genomic adaptations. Other Valsa species such as *V. malicola*, primarily found on apple trees but exhibited low-to-none infectious ability, has been reported to be pathogenic on poplar in Europe [10, 11]; *V. persoonii* mainly infect poplar (Populus spp.) but was also isolated from apple trees with canker disease[12]) and *V. sordida*, mainly infects peach or cherry (*Prunus spp*.) [10, 13, 14, 15]. It was also reported to be a causal agent of sinusitis in human who is immune-compromised by acute myeloid leukemia [16]. Due to a wide range of woody hosts and the mode of infection, Valsa canker has become a major threat to fruit production.

**Figure 1.**
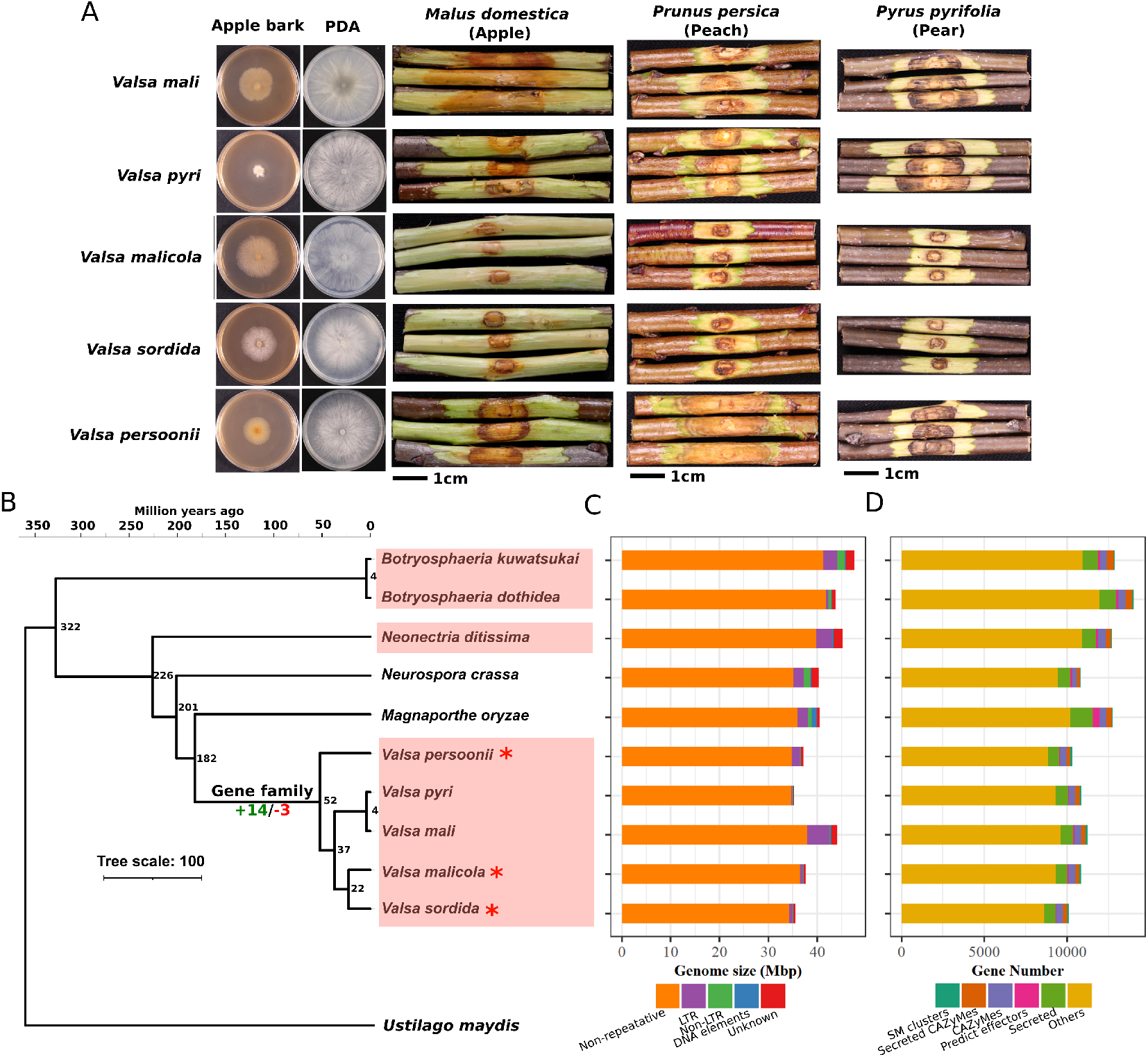
Genome and gene family evolution of five Valsa species genomes. (A) Axenic growth and woody tree branch infection by five Valsa species. Scale bars are 1cm, images were taken 7 days after inoculation. (B) Genus level phylogeny and split time estimation of five Valsa species and related fungal species. Species with * are the species sequenced in this study. Node labels are the estimated split time (million years ago). The number of gene families that expanded (14) or contracted (3) in *Valsa* clade. Branches with orange background indicate canker fungi. (C) Genome size and the proportion of different repeat sequence families including DNA elements, Non-LTR, LTR and Non-repetitive elements. (D) Number of genes annotated as secondary metabolic clusters, secreted CAZyMes, unsecreted CAZyMes, predicted effectors, secreted proteins and other functions.

In this study, we report genome assemblies and annotations for *Valsa malicola, Valsa persoonii* and *Valsa sordida*. Along with another two previously sequenced *Valsa* species *V. mali* and *V. pyri*, we conducted a comparative genomics analysis and give insights on potential roles of different genomic features in host adaptation and pathogencity development. We show that local genomic diversification are highly related to membrane transporter activities which facilitate anti-drug pumping, nutrient acquisition and anti-oxidation. Gene families of these activities are undergoing expansion in Valsa clade. Lineage specific loss or retention of genes and secretion signals are consistent with their host preference development. Extensive variations in repeat sequence contents and AT-rich regions induced by RIP are likely playing roles in secondary metabolic gene cluster diversification and integrity maintenance.

## Results

### Genome assembly and annotation of five *Valsa* species

Paired-end sequencing was performed with genome DNA samples isolated from *V. malicola, V. sordida* and *V. persooni* by an Illumina Hiseq platform. Sequencing depth for V.malicola, V.sordida and V.persooni are 147.3x, 195.6x and 154.1x respectively, genomic sequences were assembled by ABySS [17] into 287, 260 and 353 scaffolds which represent 98.9%, 98.6% and 98.9% of the whole genomes as evaluated by BUSCO [18] respectively. Proteomes predicted by MAKER suggested that V.malicola possessed the most protein encoding genes (10,848) followed by V. persooni (10,296) and V. sordida (10,099)[19]. BUSCO [18] evaluation suggested 97.9%, 98.9% and 98.6 % proteome completeness respectively. Including the previous sequenced *Valsa mali* and *Valsa pyri*, gene number comparison of functional groups of annotated genes across five species showed little to none noticeable variations except for pathogenicity related genes (Figure 1D, Table S1. Despite of the moderate variations in gene copy numbers, the five genomes showed significant size differences by ranging from 35.73 to 44.52 Mbp. This variation is largely attributable to the variations of repeat sequence contents which varied from a minimum of 2.83% for *V. pyri* genome and a maximum of 14.18% for *V. mali* genome (Figure 1 C). The higher content of repeat sequences in *V. mali* is likely due to more frequent insertion into gene bodies and secondary metabolic clusters than other four species (Figure S2, Table S2). The classification of repeat sequences in five *Valsa* species genomes based on RepeatMasker (RepBase version 20170127) and RepeatModeler (*de novo* prediction) suggested long terminal repeat(LTR) transposon elements(TEs) is the major content (Supplement Data 1). Furthermore, local GC content scanning detected a mild to moderate signal of ‘two-speed’ genome in *V. mali* but not other four suggesting a fast pace of evolution of this species (Figure S3E) Next, we used 1,250 single copy syntenic genes across 11 fungal relatives including five *Valsa* species mentioned hereinbefore, additional three fungal species causing woody three canker disease *Botryosphaeria kuwatsukai, Botryosphaeria dothidea* and *Neonectria ditissima* and two well known model filamentous fungi *Neurospora crassa* and *Magnaporthe oryzae* and lastly a basidiomycete *Ustilago maydis* as an outgroup to construct a maximum likelihood phylogeny and estimate divergence time. In consistent with previous studies [7, 20], *V*.*mali* and *V. pyri* were placed together as a clade with a divergence time of about 4 million years ago (Figure 1B). Another sister clade compassing *V*.*sordida* and *V*.*malicola* was diverged from the *V. mali* and *V. pyri* clade at about 19 million years ago and the two species within in this branch diverged at 9.8 million years ago. *V. persooni* was placed in a single branch and showed to diverge from the other two sub-clades at 28 million years ago (Figure 1B). Interestingly, the other three canker fungi *B. kuwatsukai, B. dothidea* and *N. ditissima* were placed outside of the N. crassa and *M. oryzae* clades which suggested a history of host switch from woody plants to Poaceae while some new lineages diverged from the common ancestor (Figure 1 B).

### Evolution of gene families in *Valsa* species

Annotated protein sequences for five *Valsa* species were grouped into 10,688 gene families among which 7,456 were present in five species, with the remainder being present in 1-4 species (Figure 2 A & B). Of the gene families present in all five *Valsa* species, 94% (6,980) were single copy in that species suggesting gene duplication events predominantly occurred at local level other than whole genome level. Using the phylogeny and estimated split time, CAFE [21] identified 3 gene families undergoing contraction and 14 gene families undergoing expansion in the *Valsa spp*.. The three contracted gene families are involved in sulfur compound metabolism and secondary metabolism. Interestingly, these gene families have more gene copies in *B. kuwatsukai, B. dothidea* and *N. ditissima* which represent ancestral lineages of canker fungal pathogens. Moreover, despite of a significant contraction in gene copy numbers, these gene families were not completely purged in *Valsa* clade, instead, were retained with relatively high copy numbers suggesting their indispensable roles for host colonization by *Valsa spp*. (Figure 2 C). On the other hand, the 14 gene families undergoing expansion showed an enrichment in annotations related to trans-membrane transport, secondary metabolic process and pyrophosphatase activity. These gene families might have played important roles in host expansion during the divergence of clade (Figure 2 D). Among the 13 expanding gene families, 6 of them have significantly more gene copies than their ancestral canker relatives (students t-test), in contrast to the gene families undergoing contraction, these functions might be essential for woody plant colonization while an enhancement would be favorable for ornament tree infection by *Valsa spp*. (Figure 2D).

**Figure 2.**
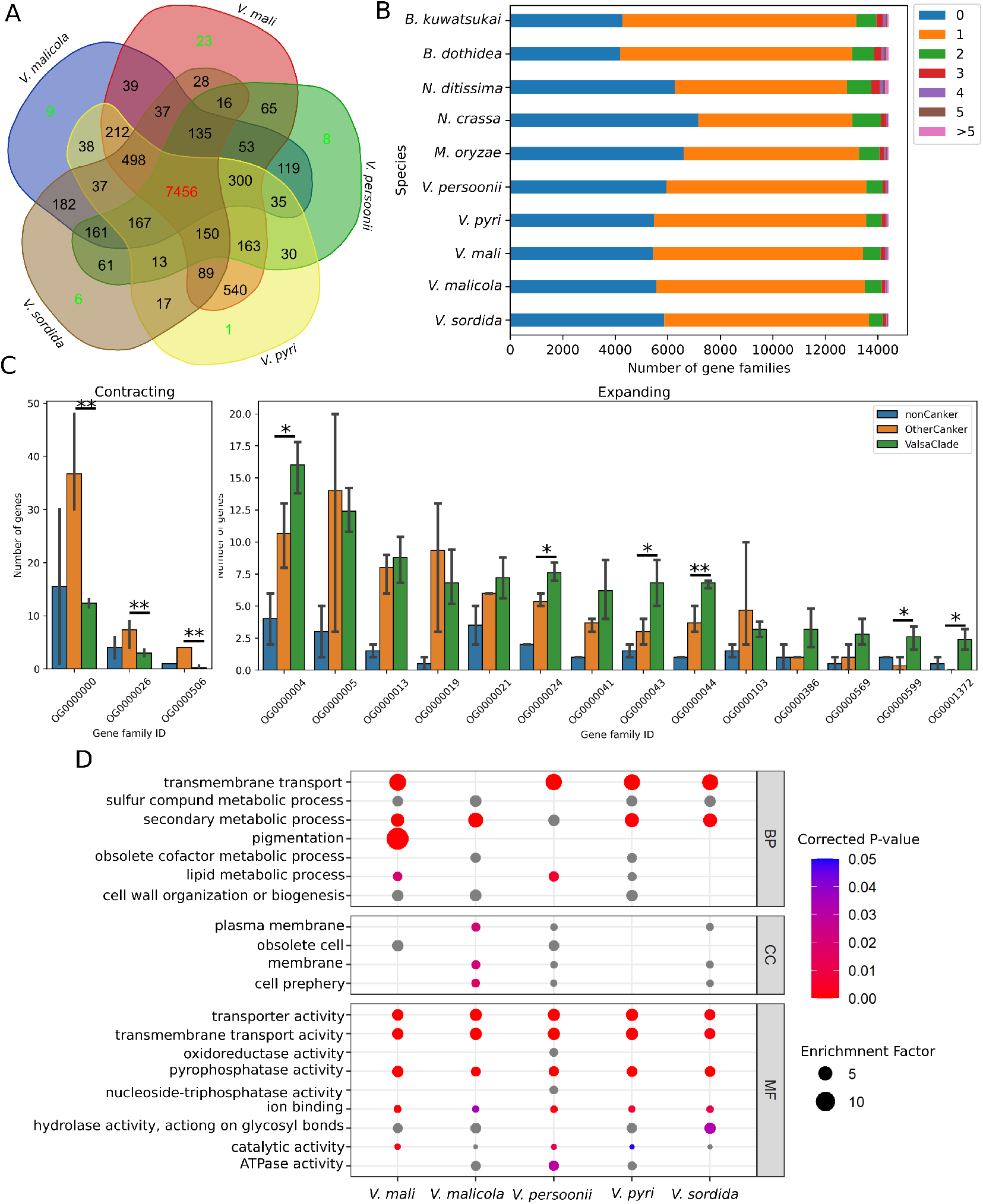
Evolution of gene families in *Valsa* species. (A) Conservation and specification of gene families across five *Valsa* genomes. (B) Gene families with different copy numbers of genes in the species included in the phylogeny. (C) Bars indicating number of genes in the Valsa clade contracted and expanded gene families across three groups of species included in the phylogeny as shown in Figure (1B) (D) Gene ontology terms overrepresented in the significantly expanded gene families in *Valsa* clade.

### Local diversification and selection in *Valsa spp*. genomes

Generally, major chromosomal rearrangements are very rare in eukaryotic genomes which lead to large collinear blocks compassing many genes across relative species. However, fungal present a very different pattern where microsyntenic regions with randomized orientation resulted from small local chromosomal rearrangements are more frequent in a mode of evolution called mesosynteny [22]. Comparing to *V. mali* genome, the genome of other four *Valsa* species retained an overall more than 80 % synteny (Figure 3A & B) while several inter- and intra-chromosomal rearrangments were also observed (Figure 3A & Figure S3 A). Most of the duplications were consistently observed in comparisons between *V. mali* and the other four species respectively. A large segmental duplication between chromosome 6 and 2 was only observed in the comparison between *V. persoonii* and *V. mali* genomes (Figure S3 A). Moreover, we identified a set of orthologous pairs that lost synteny and exhibited sequence diversification at a significantly higher level than that of the same number of randomly selected syntenic genes (Figure 3 C & D). Sequence diversification of genes underlies neofunctionalization and drives phenotypic adaptation of an organism. We therefore examined the functions of the genes with nonsynonymous mutation (Ks) value higher than 1.5 and found significant enrichment of gene ontology (GO) terms associated with membrane transporter activities such as xenophobic transmembrane transport, ion transport, organic acid transport and oligopeptide transport; protein binding activities such as modified amino acid binding and iron ion binding; plant signal response such as response to karrikin; anti-oxidation such as oxidoreductase activity, fungal-type cell wall biogenesis and secondary metabolite biosynthesis (Figure S4). Further functional classification on the set of genes associated with transmembrane transporter activities revealed that a substantial portion of which are members of Major Facilitator Superfamily (MFS), particularly Drug:H+ Antiporter (DHA) transporters which function as efflux pumps to confer resistance to anti-fungal drugs [23, 24] and oxidative stresses [25] (Figure S5). A large number of nutrient transporter families such as sugar transporter (SP) and APCs are also included in this set of genes. To explore lineage specific diversification of genes during evolution, we identified 2334 single-copy orthologues among *Valsa spp*., *Magnaporthe oryzae, Neurospora crassa* and *Ustilago maydis* using orthomcl [26] and detected selection signatures in the *Valsa* clade using the branch-site model implemented in PAML [27] (Figure S1A)(see Methods). The results showed that genes related to fatty acid beta-oxidation, DNA helicase activity and ATPase activity were over-represented in those genes with significant signals for positive selection (Figure S1B).

**Figure 3.**
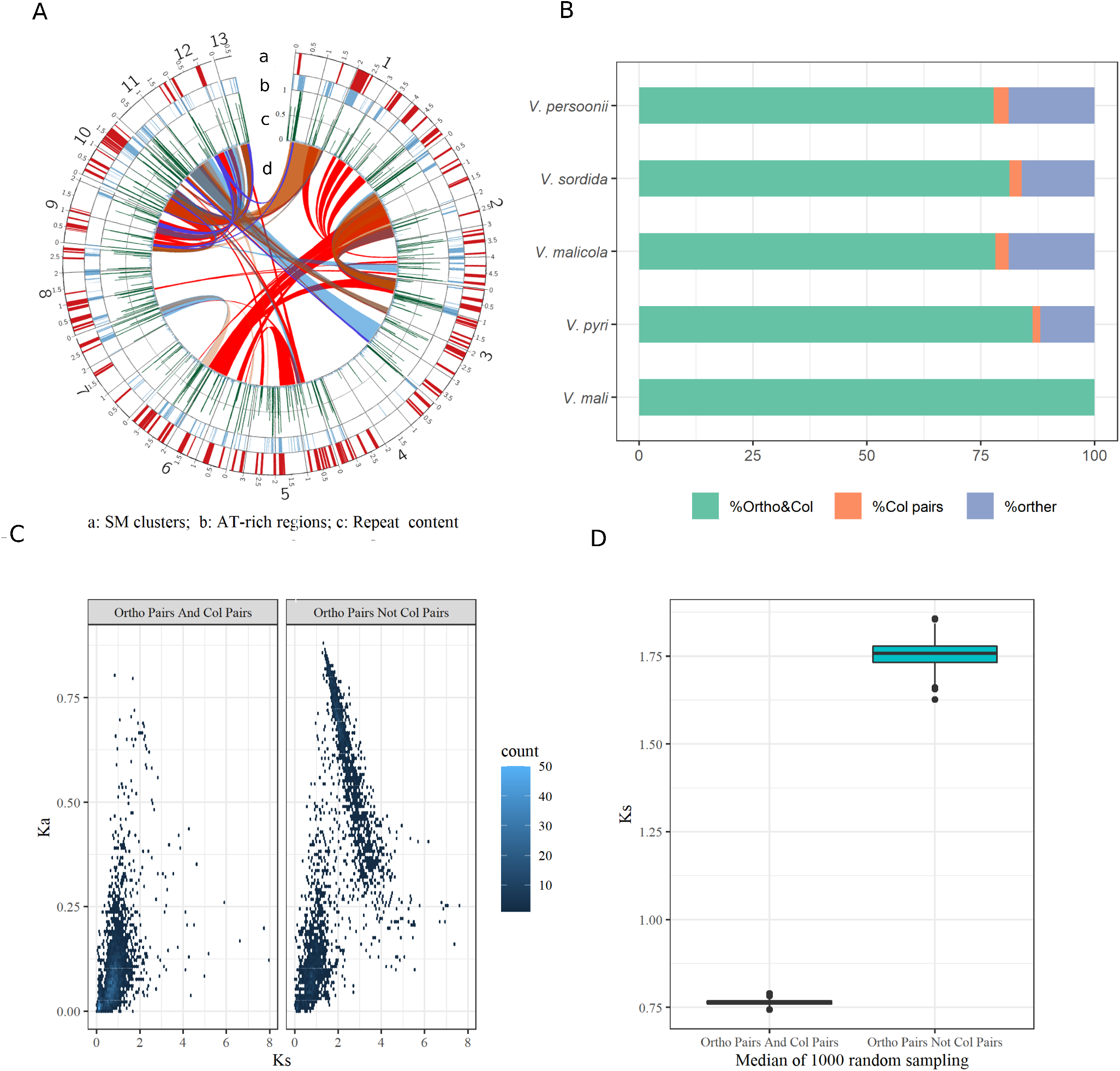
Genomic conservation and diversification across five Valsa species. (A) Circos plot showing a: Secondary metabolic gene clusters; b: AT-rich regions; c:Repeat content and d: Chromosomal local rearrangements in different species compared to *V. mali*. (B) Proportion of orthologous genes in synteny blocks and orthologues lost synteny with *V. mali*. (C) Comparison between nonsynonymous substitution (Ks) and synonymous substitution (Ka) of 3000 genes randomly selected from syntenic orthologues (left) and non-syntenic orthologues (right), respectively. (D) Comparison of distribution of median KS of 3000 randomly selected genes from syntenic orthologues and orthologues lost synteny for 100 times.

### Association between repeat sequences with lineage specific genetic variations in *Valsa* .spp

Repetitive sequences not only led to genome size variation but also increased genomic plasticity and drove host/environmental adaptation by affecting genome architecture [28, 29]. The genome size variation among five *Valsa* genomes was largely attributable to repeat content (Figure 1C). In fungal genomes, repetitive sequence and transposable element activities were contained via repeat induced C to T point mutation (RIP) to maintain genome stability leading to depletion of GC content and thus, enrichment of AT content in the genomic regions flanking repeat sequences or transposable elements[30, 31]. Therefore, AT-rich regions can be used as a proxy of repeat sequence footprint. We consistently observed significant correlations between local repeat sequence and AT-rich regions among all five *Valsa* genomes, most repeat sequences were within 200 bp from AT-rich regions while this was not observed in *S. sclerotinium* genome which possesses low repeat content (Figure S7A). In addition, GC content of AT-rich regions in *in V. mali* genome revealed a distinct distribution which features a weak but visible “two-speed genome” signal (Figure S3C). AT-rich regions in *V. mali* genome possessed 68 genes in total (8 genes per Mbp in AT-rich regions) which evinced a role of repeat sequences in *de novo* gene generation. In consistence to this observation in *V. mali* genome, the species specific genes and recent species specific paralogs are more likely in proximal to AT-rich regions in all five *Valsa* genomes (Figure S7B & C). From the chromosomal rearrangement analysis, we noticed that a significant portion of break points of chromosomal rearrangment events identified across five species were located in AT-rich regions (Figure 3A, Figure S3A, Figure 4A).

**Figure 4.**
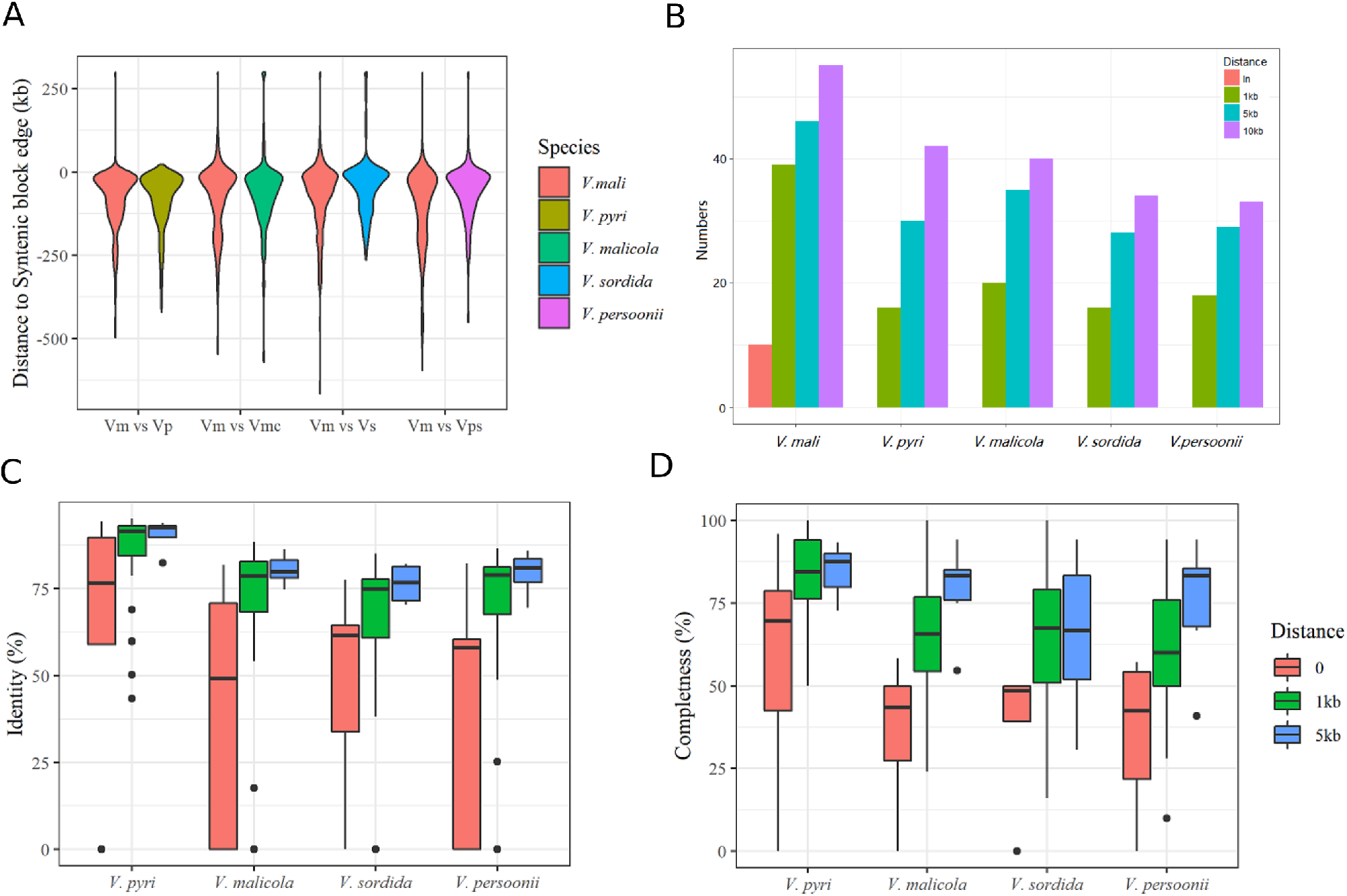
Potential role of repeat elements in SM evolution in Valsa species. (A) Position relation between AT-rich region borders and chromosomal break points (negative means break points outside of AT rich regions). (B) Number of SM detected in different distance to AT-rich regions in five *Valsa* species. (C) Relation of SM gene identities to their *V. mali* orthologues with the distance to AT-rich regions. (D) Synteny retention of SM clusters to their *V. mali* orghologues with their distance to AT-rich regions.

### Conservation and diversification of secondary metabolism clusters are associated with AT rich regions

*V*.*mali* has undergone a significant SM cluster expansion compared to fungal relatives that infect grass species [6]. This might suggest important roles of secondary metabolism for woody host by *Valsa* species. Secondary metabolites in fungi can be categorized into ribosomal peptides and amino acid-derived compounds; polyketides and fatty acid-derived compounds and terpenes [32]. The key enzymes which initiates the first biosynthetic step are classified into : Non ribosomal peptide synthetases (NRPS), polyketides synthases (PKS) and terpene synthases(TS) [32]. We predicted the types of secondary metabolic clusters identified in five *Valsa* genomes using antiSMASH with these key enzymes as queries [33] and compared the clusters across five *Valsa* species. Syntenically conserved SM clusters showed different degrees of variation in sequence and structure using *Valsa mali* as a reference (Figure S8). Terpene metabolic clusters are largely conserved while PKS clusters exhibited higher diversity. Moreover, we found that most of the secondary metabolic gene clusters were associated with AT rich regions, 3/2 of the clusters are within 5 kb away from AT rich regions (Figure 4B). Ten of the clusters in *V. mali* are even inside AT rich regions (Figure 4B). This positional proximity of SM gene clusters to AT rich regions suggested secondary clusters might have been undergoing faster sequential or structural mutations. In consistence with the notion that AT-rich regions associate with elevated mutation rates of nearby genomic regions, the level of conservation of SM gene clusters showed a strong negative correlation with the distance of to AT-rich regions (Figure 4C & D).

### Transcriptional regulation of conserved and diverged secondary metabolic gene clusters by *LaeA* during infection

It has been shown that the LaeA, a global regulator of secondary metabolism in fungi [34], controls the virulence of *Valsa mali* by regulating the transcription of a variety of secondary metabolic gene clusters that synthesize toxins and transporters that render drug resistance during infection[35]. *Valsa spp* are heterotrophic pathogens that secrete various toxins synthesized by secondary metabolic clusters to kill their host cells during the early stage of infection [36, 37]. The size and divergence of secondary metabolic clusters among the five *Valsa* genomes might be opt to colonizing different host plants in the course of evolution. However, genomic features of secondary metabolic gene clusters have limitations in interpreting the mechanisms underlying their predicted function because they are tightly controlled at the transcription level, for example, by *LaeA*[34]. We therefore sought out to investigate the association between the conservative level of SMs and their regulation by *LaeA*. A previous study identified 85 secondary metabolic gene clusters in the genome of *V. mali* [20]. To investigate the roles of these clusters for infection, we successfully knocked out the backbone genes of 40 of the clusters and reduction of pathogenicity was observed in some of them such as *VmNRPS12* [38] and *VmNRPS14* [35]. Transcriptome profiled in the invasive hypha of *V. mali* revealed that the expression of 23 of these genes were significantly down-regulated (blue shade) and 15 of them were significantly up-regulated (pink shade) (Figure 5). A previous study in *V. mali* reported the 21 of the 40 clusters shown were negatively regulated and 13 of the 40 were positively regulated by *LaeA* respectively in vegetative mycelia [35]. Interestingly, while *LaeA* was down-regulated during the infection, the genes that were positively regulated by *LaeA* in vegetative mycelia were not down-regulated during the infection and vice versa for the genes that were negatively regulated by *LaeA* (Figure 5. This observation might indicate the existence of other regulators during the infection that antagonistically control the expression of the secondary metabolism in *V. mali*. Among the five *Valsa* species, the clusters that were positively regulated by *LaeA* in *V. mali* showed higher *V. mali* specific conservation than the ones that are negatively regulated (Figure 5). Among the 13 backbone genes that were positively regulated by *LaeA* in vegetative mycelia, only orthologues of NRPS4, NRPS51 and NRPS10 were identified in all other four species. In contrast, among the 21 backbone genes that are negatively regulated by *LaeA* in *V. mali* vegetative mycelia, orthologues of 6 backbone genes including NRPS59, PKS39, NRPS25a, NRPS25, NRPS12 and NRPS18 were identified in all other four species.

**Figure 5.**
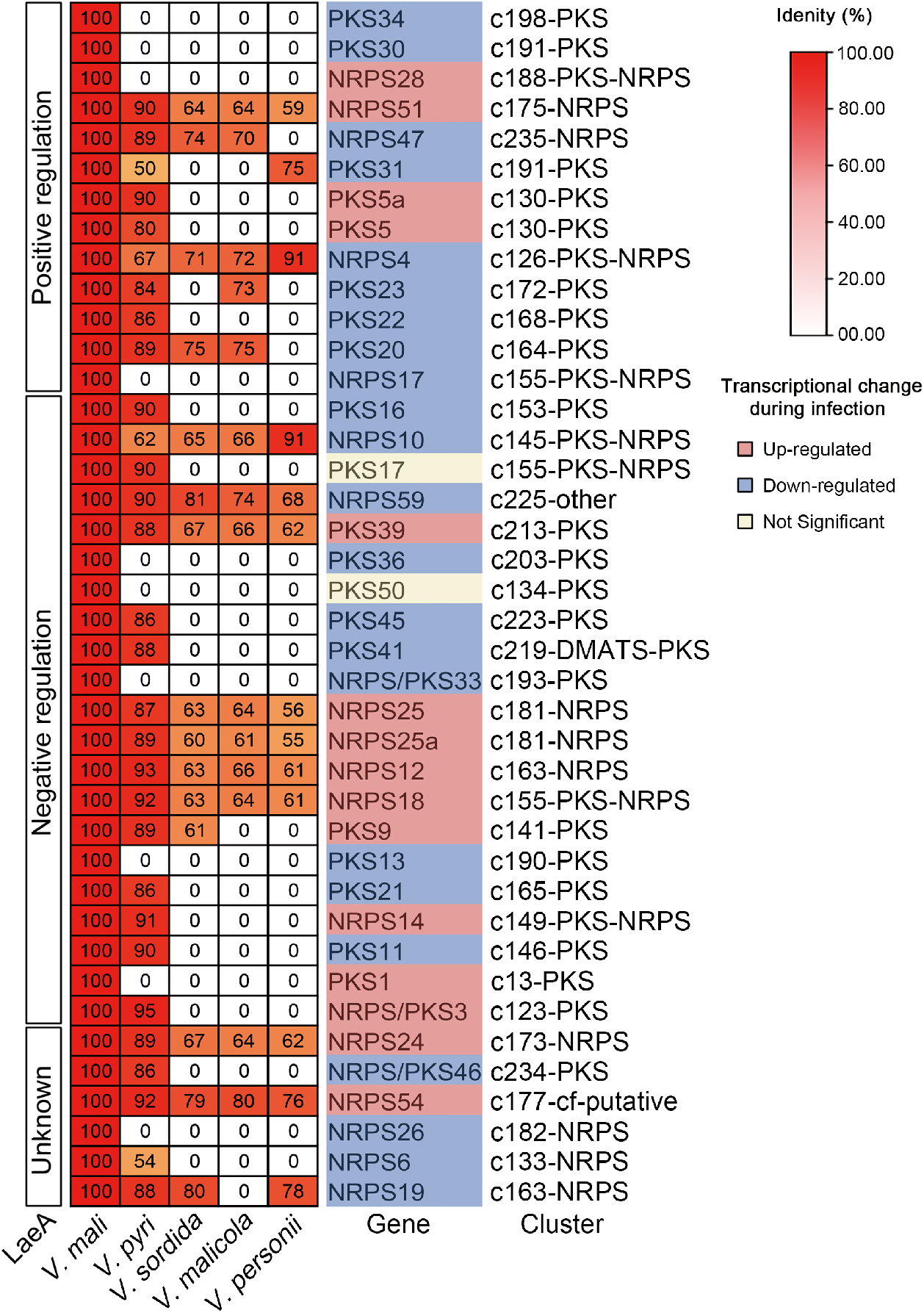
Transcriptional regulation of SM clusters by the global regulator LaeA in Valsa species. The columns in the heatmap of *V. malicola, V. persoonii, V. pyri*, and *V. sordida* show the homology similarity with *V*.*mali* secondary metabolic backbone genes, respectively; The “*LaeA*” column indicate transcriptional regulation of the SM backbone genes by the secondary metabolism global regulator *LaeA*. The “Gene” column indicate the backbone genes of the secondary metabolic gene clusters compared across the species and the shades of the gene names indicate transcriptional change during the infection. The cluster column indicates the secondary metabolic gene clusters that the backbone genes are located in.

## Discussion

*Valsa spp*. is a group of destructive pathogens on woody plants, tremendous losses are caused due to its capability of infecting a variety of important woody hosts such as apple, pear, peach and other economically important trees. The genome of *V. mali* revealed several features that facilitate its outstanding capability of infection such as expanded pectinase families, repeat sequence insertions, toxin biosynthesis and as well as effectors attained via horizontal gene transfer from its host [6]. The genome plasticity as a result of intensive genetic engagements with their host and co-exist microbe communities have risen as a promising weapon to excel local adaptation [39]. In this study, genomes of three more fungal pathogens that vary in host specificity and pathogenicity from Valsa clade (*V. persoonii, V. malicola* and *V. sordida*) were sequenced. We show that host adaptation by *Valsa* species might be driven by local genomic diversification, gene family contraction and expansion and potentially repeat sequence mediated secondary metabolic gene cluster evolution.

Comparison of the five Valsa spp genomes revealed well conserved synteny with a few segmental rearrangments and duplications (Figure S3). We have identified some intra-chromosomal inversions featuring mesosytenies as revealed in the class Dothideomycetes suggesting specific transposition of genetic elements might also be one of the mechanisms for Valsa species to increase genetic diversity [40]. However, these results need to be interpreted with cautions because among the five species, only the genome of *V. mali* was assembled and annotated to the chromosome level. The other four were only assembled to contigs which might give rise to falsely high number of inversions. The small number of chromosomal translocations observed was likely due the same reason. As was suggested in ascomycota fungi, the frequent observation of microsynteny between ascomycota distant relatives might be risen from inversion but extremely rare translocations [22], we hypothesize that *Valsa* species might not be an exemption because we did not observe frequent translocation between long contigs that are in synteny with *V. mali* chromosomes. On the other hand, the genes in syntenic blocks are losing the homology suggesting there might be species specific sequence differentiation.

Frequent local sequence diversification were enriched in the regions where the genes are largely involved in transmembrane transporter activities especially DHA transporters and respond to plant chemical compounds such as karrikin. DHA transporters are likely responsible for fungicide resistance development in fungal pathogens [41]. This genes or processes can be potential new targets for disease control. Karrikin is a group of chemical compound involved in plant seed germination and early development, it has been shown to be involved in phytohormone signaling networks [42]. Current understandings on karrikins are primarily in plant growth, therefore, it would be interesting to further investigate whether this response to karrikin is also involved in disease defense [42, 43]. Interestingly, gene loss events are usually associated with transporter and secondary metabolism in *Valsa* species but they never loss the function suggesting gene loss by *Valsa* species are more of a redundancy reduction rather than purge of certain function. This selective but balanced gene gain and loss towards certain functions in *Valsa* species might be an important mechanism for specific environment adaptation [44, 45, 46].

Repeat elements have been shown to play critical roles in fungal evolution. For example, varieties of DNA segments duplicated by TEs during their frequent translocations along the genome potentially increased the number of effector protein encoding genes and their sequence divergence in wheat stripe rust fungus *Puccinia striiformis* [47]. In cotton wilt fungus Verticillium dahliea, TE elements caused erroneous double-strand repair that mediated genomic rearrangement resulting in highly variable lineage-specific regions where active TEs are enriched and provide local genetic plasticity [48]. Based on these previous evidence, the significant variation in repeat content in *V. mali* and *V. pyri* genomes very likely promoted *de novo* genetic variation for host adaptation. In this study, we show evidence of an intimacy between repeat sequence (represented by AT-rich regions) and secondary metabolic gene cluster. Secondary metabolic clusters are critical for nutrient acquisition, infectious structure (appressorium, haustoria) formation and host infection [49, 50]. In *Valsa mali*, secondary metabolism has been implied to be very important for woody host adaptation [6]. In this study, we revealed that AT-rich regions induced via RIP TEs were particularly active in filamentous fungi which was implied to function to stabilize genetic information by mediating chromatin circularization [48]. AT-rich elements also appear in the flanking regions of gene clusters with an above-random level frequency suggesting a potential specific association with metabolic gene clusters. The function of this association might be consistent with TEs, which were occasionally identified at the flanks of gene clusters playing a role in MGC assembly and/or movement [51, 52]

Fungal SMs can be divided into four main chemical classes: polyketides, terpenoids, shikimic acid derived compounds, and non-ribosomal peptides [53]. The small molecules such as fungal toxins, hormones and other metabolites are revealed as important for fungal pathogen and even symptomless fungal endophytes for interaction with their plant hosts [54]. Some fungal pathogens even produces SMs to mimick the ones produced by plant host to mediate plant growth during the infection [55]. For example, in some cases fungal pathogens produce and secrete terpenes to manipulate host plant growth and development to support fungal growth. Conversely, it is also widely utilized by plant host as an antifungal molecule to defend infection[56]. Polyketide synthase (PKS) and Nonribosomal peptide synthase(NRPS) have been reported to catalyze biosynthesis of secondary metabolites that facilitate infection with plant hosts by fungal pathogens[57]. The mechanism of cluster formation and integrity maintenance are poorly understood.

In animal or allopolyploidy plant systems, micro- or mini-satellites were reported to be involved in chromosome break point formation which eventually lead to chromosomal rearrangments [58, 59]. Our observation of chromosomal rearrangment break points being frequently located in SM flanking AT-rich regions might indicate similar mechanism in fungi that at the current stage of evolution, the AT-rich regions induced by repeat sequences likely played a protective role to avoid beak point genesis from disrupting SM clusters. Furthermore, *Valsa mali* has been shown to be heterothallic thus sexual reproduction occasionally occurs in nature [60]. Recombination during sexual reproduction is a major source to gain genomic diversity, if recombination happened in matured SM clusters, the function of the SM clusters might be disrupted. In Microbotryum species, significant low recombination rates in region enriched in transposable element contents in mating-type chromosomes was observed [61]. Furthermore, recombination hotspots were found to have higher G/C repeats [62], AT-rich regions might act as a dilution of recombination hotspots as a protective mechanism. These together could explain the high synteny conservation of the clusters that are 1 to 5 kb away from AT-rich regions.

The regulation of secondary metabolic clusters are very complex, *LaeA* acts as a global regulator of the expression of SM backbone genes during the infection [34, 35]. The clusters that are regulated by *LaeA* in *Valsa mali* are not always present in other four *Valsa* species perhaps as the result of expansion of SM clusters in *V. mali*(Figure 5). *V. malicola* and *V. sordida* exhibited lower virulence compared to other tested *Valsa* species (Figure 1A) and showed the low similarity of secondary metabolic clusters with *V. mali* (Figure 5). Interestingly, *V. persoonii* which exhibited strong pathogenicity on all three hosts tested (apple, peach and pear) showed the lowest similarity of secondary metabolic cluster with *V. mali*. This might indicate even though expansion of secondary metabolism in *V. mali* contributed to its high virulence and drug resistance [20], it might not be a deterministic factor to virulence for *Valsa* species. Furthermore, the expression level change of the SM cluster backbone genes did not correspond to the supposed pattern when *LaeA* is down-regulated during the infection. For example, NRPS59, PKS39, PKS36, PKS50, PKS45, PKS41 and NRPS/PKS33 were up-regulated as expected during the infection as *LaeA* down-regulation would de-represss the expression of these genes (Figure 5). However, there was a number of genes that were negatively regulated by *LaeA* that were down-regulated during the infection (Figure 5) suggesting existence of other regulators during the infection or temporal regulation by *LaeA*. Future studies including more closely related *Valsa* species with more complete genome assembly such as long read sequencing that can resolve repeat regions would further advance our understandings on the roles of repeat sequences in pathogenicity evolution.

## Materials and methods

### Phylogeny based on whole genome single copy orthologues

Single copy proteoms were aligned by mafft using –auto mode. The alignments were then trimmed by Gblocks “Gblocks GblockAlignPath -t=p -b1=5 -b2=6 -b3=8 -b4=10 -a=y”. So the conserved blocks identified by Gblocks were concatenated. Phylogeny with brach length was constructed by iqtree with command: “iqtree -s GblockAlignPath-gb.seq -o moryz_MGG_13474T0,ncras_EAA35109 -nt AUTO -m MFP -bb 1000”. A total of 5 *Valsa* species were used in this study, namely *Valsa mali* isolate 03-8 and *Valsa malicola* isolate 03-1 isolated from apple trees, and *Valsa pyri* isolate isolated from pear trees. SXYL134, a *Valsa persoonii* (= leucostoma) isolate SXYLt isolated from a peach tree, and a *Valsa sordida* isolate YSFL isolated from a poplar tree. Among them, the genomes of *V. mali* and *V. pyri* have been sequenced and assembled [6], and the genomes of three other were sequenced in this study. All of the above strains were cultured and preserved on PDA medium according to conventional methods, and provided by the State Key Laboratory of Crop Stress Biology in Arid Regions of Northwest A & F University. For the convenience of description, five strains are referred to as *V. mali*, Vm; *V. pyri*, Vp; *V. malicola*, Vmc; *V. persoonii*, Vps; *V. sordida*, Vs. For previous transcriptome data of Vm was received from the previous article [6]

### Branch inoculation assay

Peach, pear, and apple tree branches with consistent thickness were cut into short sections with 10 cm in length. After rinsing with tap water, the branch surfaces were soaked in 6 % sodium hypochlorite solution for 15 minutes and rinsed by sterile tap water for three times. The ends of the branches were then sealed by paraffin wax after air drying. Each of the branches were burnt wounded by a soldering gun with a 5 mm punch. Colonies of each of the *Valsa* species revived for two consecutive times were cultured in PDA media and were incubated in the dark at 25 ^*°*^*C* for 48h before they were inoculated to the wounds carefully. Sterile PDA media punches were inoculated to the wounds as negative controls. The inoculated branches were incubated in containers that kept high internal moisture at 25 ^*°*^*C* for 7 days. Lesion lengths were then measured for each of the branches inoculated with each of the fungal cultures. Five wounded branches of each of the three tree species were inoculated for each *Valsa* species and the experiments were conducted in triplicates. Test the difference in pathogenicity of 5 decay fungi on apple tree branches. Take healthy branches of the apple orchard, cut the branches to about 30cm, and inoculate the branches after the decomposing bacteria are activated by PDA culture. Each test was inoculated with 3 shoots of each strain and repeated 3 times.

### Genome sequencing, assembly and gene predection

Illumina HiSeq sequencing technology was used to sequence the whole genome of three strains, the sequencing depth was Vmc 147.3x, Vps 154.1x, Vs 195.6x. After sequencing the paired-end reads, we used ABySS version 1.9.0 [17] to perform de novo assembly of the genome, and used Benchmarking Universal Single-Copy Orthologs (BUSCO) 2.0 [18] for assessing assembly completeness. The protein-coding genes were annotated using MAKER version 2.31.8 [19], and the predicted completeness of the predicted proteome was also verified using BUSCO 2.0.

### Repeat Sequences and AT-rich regions

Combined with REPETMODELER and REPEATMASKER [63], the repeat sequences of 5 strains were detected. First, REPEATMODELER was used to perform de novo prediction of the repeat sequences. The obtained consensus.fa file was integrated with the repeat sequence database of the fungus in RepBase [64], and then the integrated database was used as the input of REPEATMASKER for repeat sequences prediction of the five species. The repeat type and content ratio were calculated using a self-written Perl script. OcculterCut version 1.1 [31] was used to analyze the AT-rich regions of five strains, and the positional relationship between genes and repeats and AT-rich regions was calculated based on the analysis results.

### Genome Annotation

In order to facilitate subsequent comparative analysis, the same method was used to annotate different functional categories for the five strains. The National Center for Biotechnology Information (NCBI) online BLAST service [65] was used for NR database annotation. The Perl script, iprscan5.pl, annotated the InterPro database ([66]), and then uses Blast2GO version 4.0 [67] to combine the results to obtain GO function annotations. The Kyoto Encyclopedia of Genes and Genomes (KEGG) [68] provides an online service (https://www.kegg.jp/blastkoala/) for annotation of the KEGG metabolic pathway. Pfam 30.0 [69] database was used to annotate protein domains through pfamscan.pl. Pathogen host interactions (PHI) [70] database provides an online service (http://phi-blast.phi-base.org) to find potential pathogen-related proteins in rot fungi. SignalP version 4.1 [71] (http://www.cbs.dtu.dk/services/SignalP/) was used to predict the secretion signals of proteins, TMHMM version 2.0 [72] (http://www.cbs.dtu.dk/services/TMHMM-2.0/) predicts the transmembrane region of the protein sequence, in which the presence of a secretion signal is predicted by SingalP and the protein of the transmembrane region does not exist after 30 amino acids acting as a potential secreted protein. The predicted secreted proteins were feed to EffectorP version 2.0 [73] to predict potential effector proteins. Annotations of cell wall degrading enzymes were retrieved by hmmscan using the CAZymes HMM model file provided by dbCAN [74], and the results were classified and counted by the hmmscan-parser.sh scrip. Blastp was used to search for MEROPS [75] and Transporter Collection Database (TCDB) [76] to identify proteases and membrane transporters, respectively. Hmmscan was used to search the Lipase database (http://www.led.uni-stuttgart.de/) to identify lipases. Secondary metabolic gene clusters were predicted using the antiSMASH version 4.2 [33] tool. The predicted gene clusters were combined with InterPro annotation results to use the online tool Secondary Metabolites by InterProScan (SMIPS, https://sbi.hki-jena.de/smips/index.php) [77] to identify secondary metabolite synthesis backbone genes.

### Gene family clustering analysis

BLASTP was performed for an all vs all alignment of the proteomes of the five species, and then OrthoMCL version 2.0.9 [26] was used to find homologous gene pairs, where the parameters were set with the sequence identity threshold as 50% and e value as 1e ^*−*^5. Then use mcl to cluster OrthoMCL results, where the inflation value is set to 1.5. In order to clarify the phylogenetic relationship between the five strains, other six fungal species were introduced in, and 2213 single-copy orthologous gene families were obtained using OrthoFinder [78]. MAFFT version 7.402 [79] was used for aligning these gene families. The tool linsi in the multi-sequence alignment was used. After concatenating all the alignment results, Gblocks version 0.91b [80] was used to extract 794987 conserved amino acid sites for phylogenetic tree construction. The phylogenetic tree was inferring by IQ-TREE [81] with parameters “-m MFP -bb 1000”.

### Synteny analysis

Synteny Mapping and Analysis Program (Symap) version 4.2 [82] was used to conduct pairwise genomic collinearity. Contigs shorter than 10 Kbp were filtered out, and gene pairs within collinear blocks were identified using MCScanX [83]. Because *Valsa mali* (Vm) are the only species with complete gensequenced and there are many gene fragments of other strains, Vp is used as a reference when analyzing genomic sequence recombination. There are two cases, 1) trans-chromosomal recombination, that is, multiple chromosomes of Vm correspond to a single genomic sequence of others, mainly considering transposition of synteny blocks and inversion of individual genes in the block, 2) inversion in chromosomes, single chromosome of Vm corresponds to a single sequence of other species, considering the transposition and inversion of blocks, and the insertion and deletion of sequences between adjacent blocks.

### Branch site positive selection analysis

ParaAT.pl [84] was used to transform the amino acid multiple sequence alignment results obtained from the MAFFT alignment into codon alignment. Single-copy orthologous genome families among 7 fungi (5 *Valsa* species, *Magnaporthe oryae* and *Sclerotinia sclerotiorum*) were selected for analysis using the codeML tool in PAML [27]. CodeML’s brach-site model (model = 2, Nsites = 2) uses two hypotheses to analyze each branch separately, where the null hypothesis (H0) parameter is set to fix_kappa = 0, fix_omega = 1, alternative hypothesis (H1) The parameters are set to fix_kappa = 0 and fix_omega = 0. The two hypotheses obtain the likelihood values LnL0 and LnL1, respectively, and calculate the P value of 2 x (LnL1-LnL0) by the chi-square test, and then correct the P value using the FDR (Benjamini-Hochberg Procedure) method. Finally, the gene family under the role of positive selection was determined, requiring a corrected P value of less than 0.05, and positive selection sites were detected by Bayes Empirical Bayes (BEB) in the alternative hypothesis result. For positive selection between gene pairs, KaKs_Calculator version 2.0 (Wang 2010) was used for analysis. Considering that the positive selections that can be detected are generally at a few positions, the split.class tool provided by KaKs_Calculator was used to segment the sequence alignment result. The segmentation method is a 57 bp base codon box. The shift length is 6 bp. Finally, KaKs ratio >1 and fisher test p value less than 0.05 were considered to have positive selection.

### Transcriptional regulation of secondary metabolic gene clusters across five *Valsa spp*

To investigate transcriptomic changes of the secondary metabolic gene clusters during the infection of apple branch by Valsa mali. We isolated RNA samples from the invasive hypha of *V. mali* from the colonized apple tree branches and performed RNA sequencing using an Illumina sequenser (Illumina, San Diego, CA, USA). Because 40 of the 85 secondary metabolic gene clusters predicted in the genome of *V. mali* [20] have been successfully knocked out. We therefore focused on these 40 SM cluster backbone genes. TBtools [85] was used to fetch the coding sequences (cds) of 40 backbone genes of secondary metabolic gene clusters that were knocked out from *V. mali*. The CDS fetched by TBtools were then translated into amino acid sequences and used as queries to search for homologes in *V. malicola, V. persoonii, V. pyri* and *V. sordida* locally constructed protein database by blastp [65, 86]. The hits with E-value lower than 1e^-10^ and identity higher than 50% were considered as orthologues.

## Acknowledgements

This work is funded by the National Natural Science Foundation-Xinjiang Joint Foundation of China (U19032061007919). We thank Professor Chengmin Shi from Hebei Agricultural University for the constructive suggestions on the layout of this study which greatly improved the structure and outcome of this study.

## Author contribution

L. H., G. S., S. X., and M. Z., conceived this research, S. X., conducted genome assembly and annotation, S. X., G. S., and C. Z., conducted the bioinformatics analysis, L. T., performed the secondary transcriptomics comparison across five species, G. S., S. X., and L. H., drafted the manuscript. All authors approved the final version of the manuscript.

## Data availability statement

Genome assembly and annotation of *Valsa mali* were retrieved from NCBI with assembly ID: ASM81815v1 under BioProject: PRJNA268126; Genome assembly and annotation of *Valsa pyri* were retrieved from NCBI with Assembly ID: ASM81338v1 under BioProject: PRJNA296468; Genome assembly and annotation of *Valsa malicola* can be accessed from NCBI with Assembly ID: ASM379531v1 under BioProject: PRJNA296468; Genome assembly and annotation of *Valsa sordida* can be accessed from NCBI with Assembly ID: ASM379527v1 under BioProject: PRJNA296468; Genome assembly and annotation of *Valsa persoonii* can be accessed from NCBI with Assembly ID: ASM379529v1 under BioProject: PRJNA296468. All of the data associated with the figures are included in this manuscript.

---

**Table S1.** Number of pathogenicity related genes

|  | SM clusters | Predict effector | CAZyMes | Secreted CAZyMes | Secreted | Others |
| --- | --- | --- | --- | --- | --- | --- |
| *Valsa mali* | 127 | 79 | 400 | 254 | 754 | 9604 |
| *Valsa persoonii* | 107 | 60 | 391 | 224 | 663 | 8851 |
| *Valsa malicola* | 114 | 75 | 414 | 237 | 695 | 9313 |
| *Valsa pyri* | 114 | 65 | 374 | 262 | 738 | 9302 |
| *Valsa sordida* | 100 | 69 | 386 | 257 | 679 | 8608 |
| *Botryosphaeria dothidea* | 70 | 145 | 423 | 432 | 995 | 11945 |
| *Botryosphaeria kuwatsukai* | 63 | 132 | 406 | 414 | 930 | 10927 |
| *Magnaporthe oryzae* | 53 | 428 | 390 | 348 | 1334 | 10202 |
| *Neurospora crassa* | 19 | 107 | 297 | 206 | 750 | 9434 |
| *Neonectria ditissima* | 43 | 113 | 489 | 301 | 845 | 10894 |

**Table S2.** Repeat sequence insertion in gene bodies and secondary metabolic gene clusters

|  | In genes |  | In SM clusters |  |
| --- | --- | --- | --- | --- |
|  | Number | Length(kb) | Number | Length(kb) |
| *Valsa mali* | 111 | 67.1 | 40 | 648.2 |
| *Valsa pyri* | 74 | 26.2 | 1 | 4.1 |
| *Valsa malicola* | 98 | 53.1 | 19 | 113.3 |
| *Valsa sordida* | 50 | 19.1 | 6 | 36.0 |
| *Valsa persoonii* | 59 | 23.0 | 3 | 23.0 |

**Table S3.** Annotation of species specific genes

|  | SM localization | Pfam | PHI | Secreted | Total |
| --- | --- | --- | --- | --- | --- |
| V. mali | 254(7.25E-05) | 188 | 121 | 62 | 974 |
| V. pyry | 127(0.5203) | 91 | 66 | 41 | 738 |
| V. malicola | 196(6.4E-04) | 234 | 159 | 67 | 969 |
| V. sordida | 157(3.48E-06) | 147 | 117 | 50 | 705 |
| V. persoonii | 248(1.60E-10) | 280 | 175 | 97 | 995 |

**Table S4.** GO terms enriched in species specific genes

|  | V. mali | V. pyry | V. malicola | V. sordida | V. persoonii |
| --- | --- | --- | --- | --- | --- |
| tetrapyroole binding | 1.98E-08 | 1.76E-02 | 1.67E-06 | 6.96E-06 | 1.75E-11 |
| cofactor binding | 2.84E-08 | 5.08E-02 | 4.45E-06 | 4.17E-06 | 3.98E-07 |
| heme binding | 2.93E-08 | 1.87E-02 | 5.43E-06 | 6.30E-06 | 1.84E-11 |
| iron ion binding | 1.38E-06 | 2.39E-02 | 3.70E-06 | 3.91E-05 | 1.01E-08 |
| oxidoreductase activity | 2.09E-05 | 2.31E-02 | 1.20E-05 | 2.19E-09 | 1.78E-11 |

**Figure S1.**
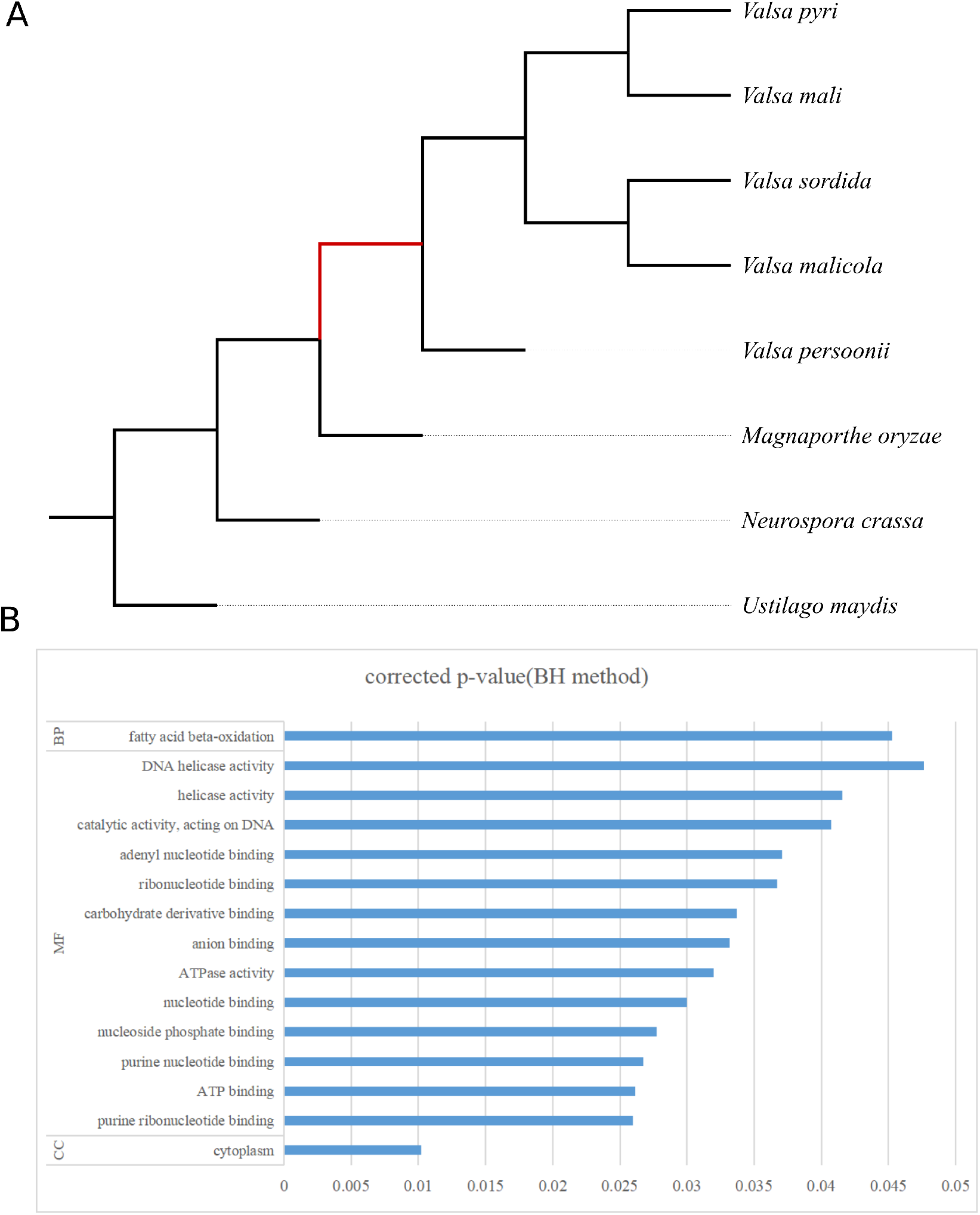
Orthologous genes with significant selection signal in Valsa spp. branch. (A) The phylogeny used for orthologous gene pair kaks analysis. The clade labeled in red indicates the foreground clade used as targets in the codeml implemented in PAML[27], *Magnaporthe oryzae, Neurospora crassa* and *Ustiliago maydis* were used as background. (B) Gene ontology terms over-represented in the gene sets with positive selection signals in Valsa clade.

**Figure S2.**
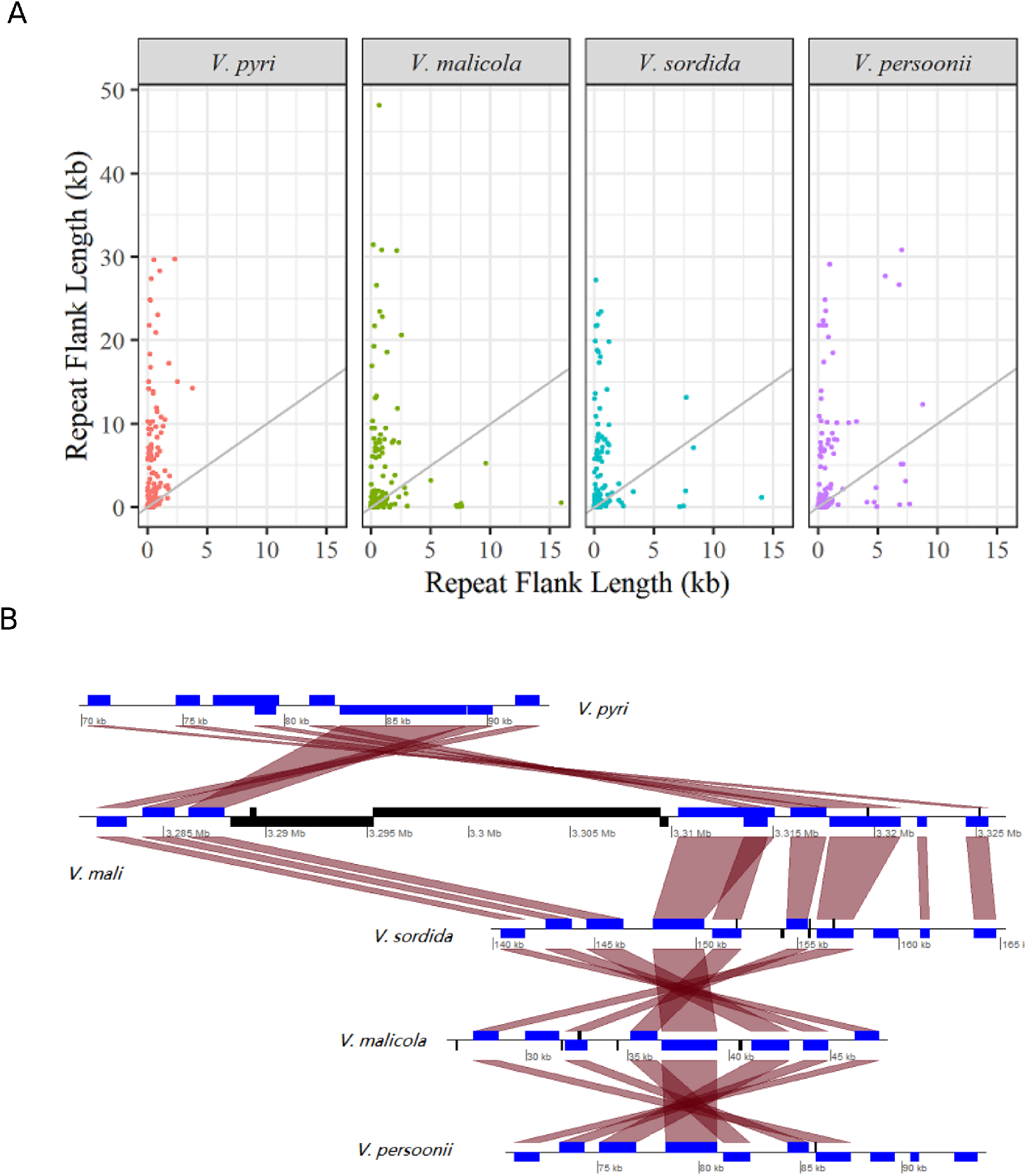
Repeats expansion in synteny blocks of V. mali. (A) Jetterplot shows the repeat length difference in each synteny blocks between *V. mali* and other four valsa species. (B) Long repeats insert into *V. mali*. The blue blocks are genes, the black blocks are inserted repeats.

**Figure S3.**
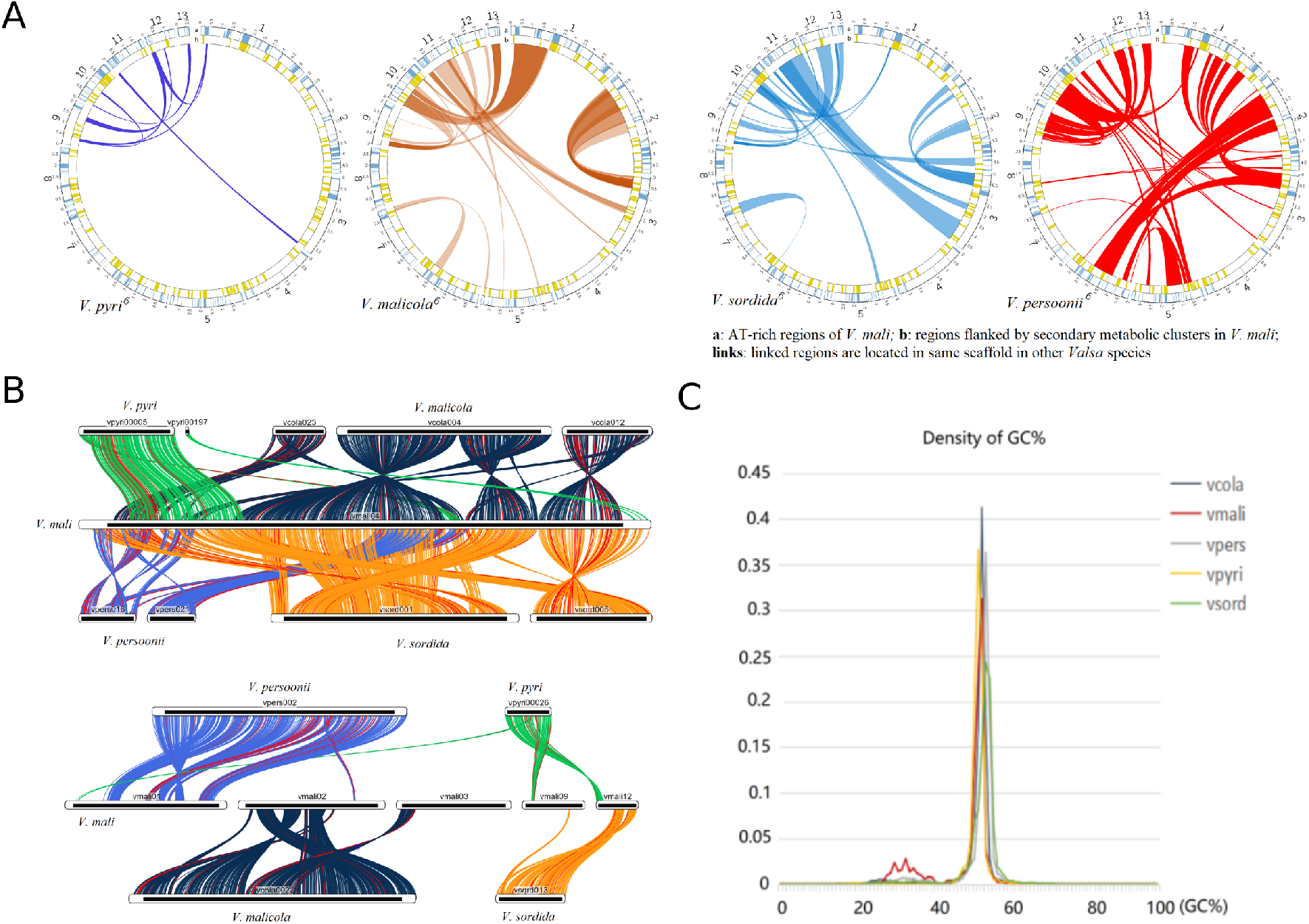
Local chromosomal rearrangments of Valsa genomes. (A) Inter- and intra-chromosomal rearrangement events identified among four species using *V. mali* as a reference; (B) Examples of Inter- and intra-chromosomal rearragnment events; (C) A mild ‘two-speed’ genome signal detected in *V. mali* genome

**Figure S4.**
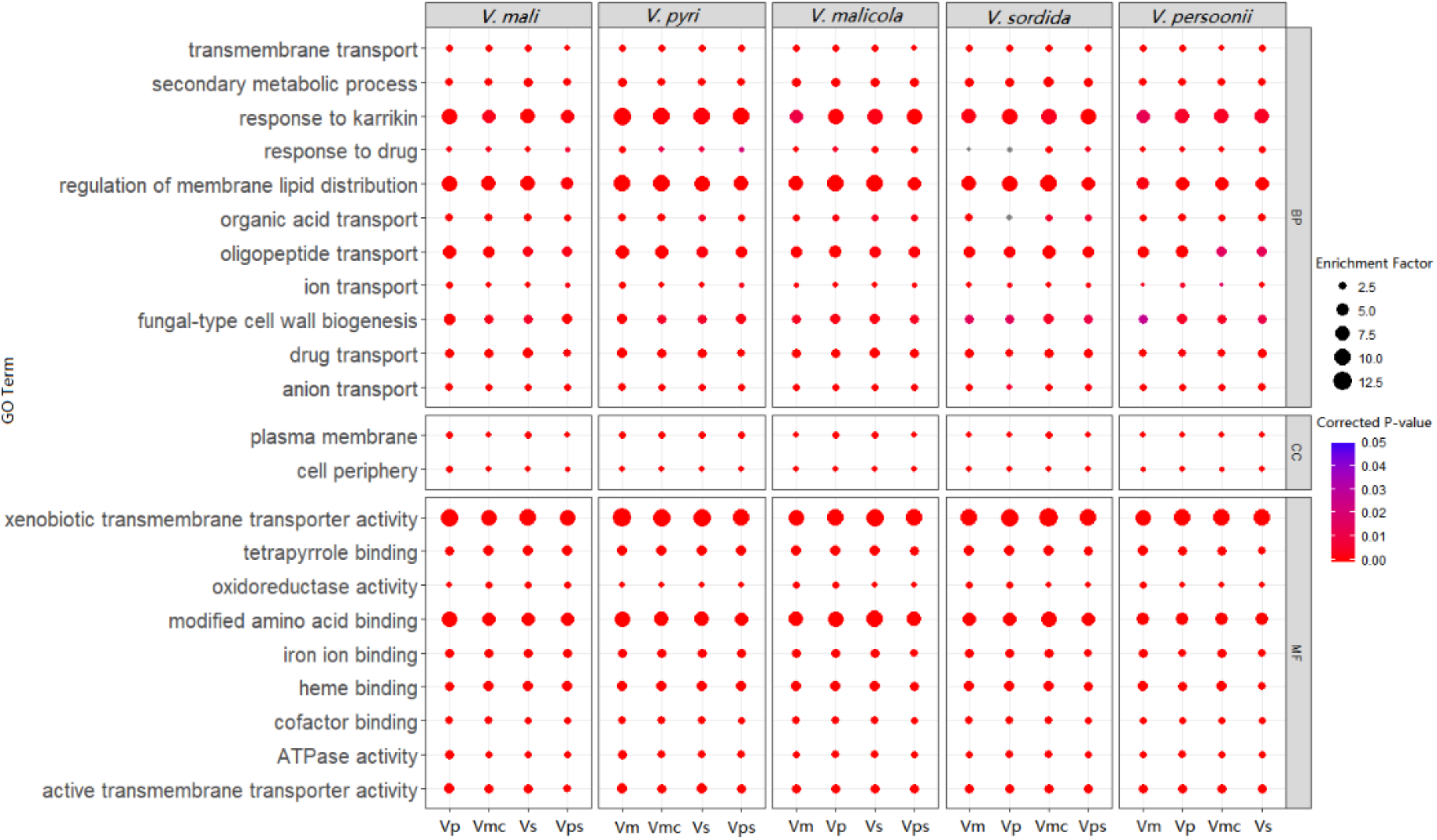
Gene ontology terms over-represented in orthologous genes lost synteny that under selection. (A) Gene ontology terms enriched, size indicate enrichment factor, corrected p values are color coded.

**Figure S5.**
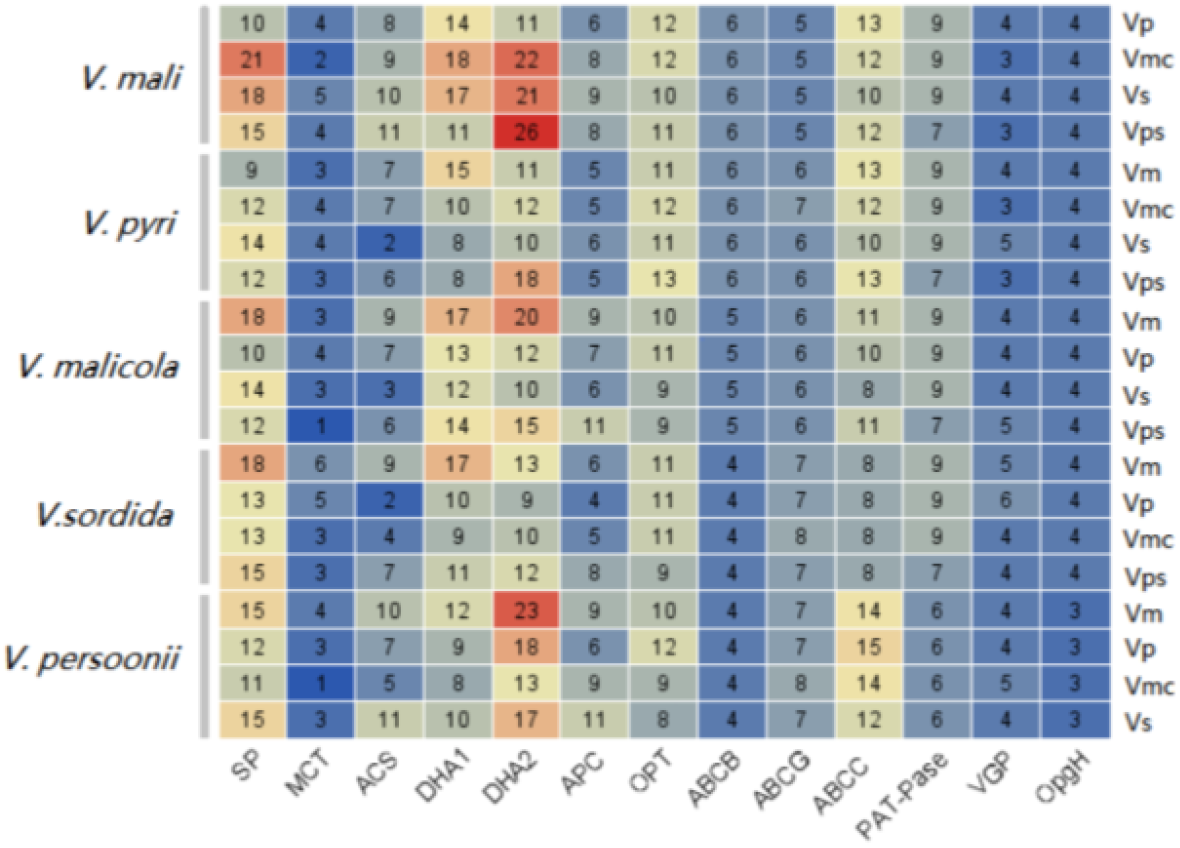
Membrane transporter family enriched in highly dynamic (KS >1.5) orthologues that lost synteny in Valsa species. (A) Number are gene counts among each transporter family and color in each squre indicate proportion of the counted genes in same transporter family

**Figure S6.**
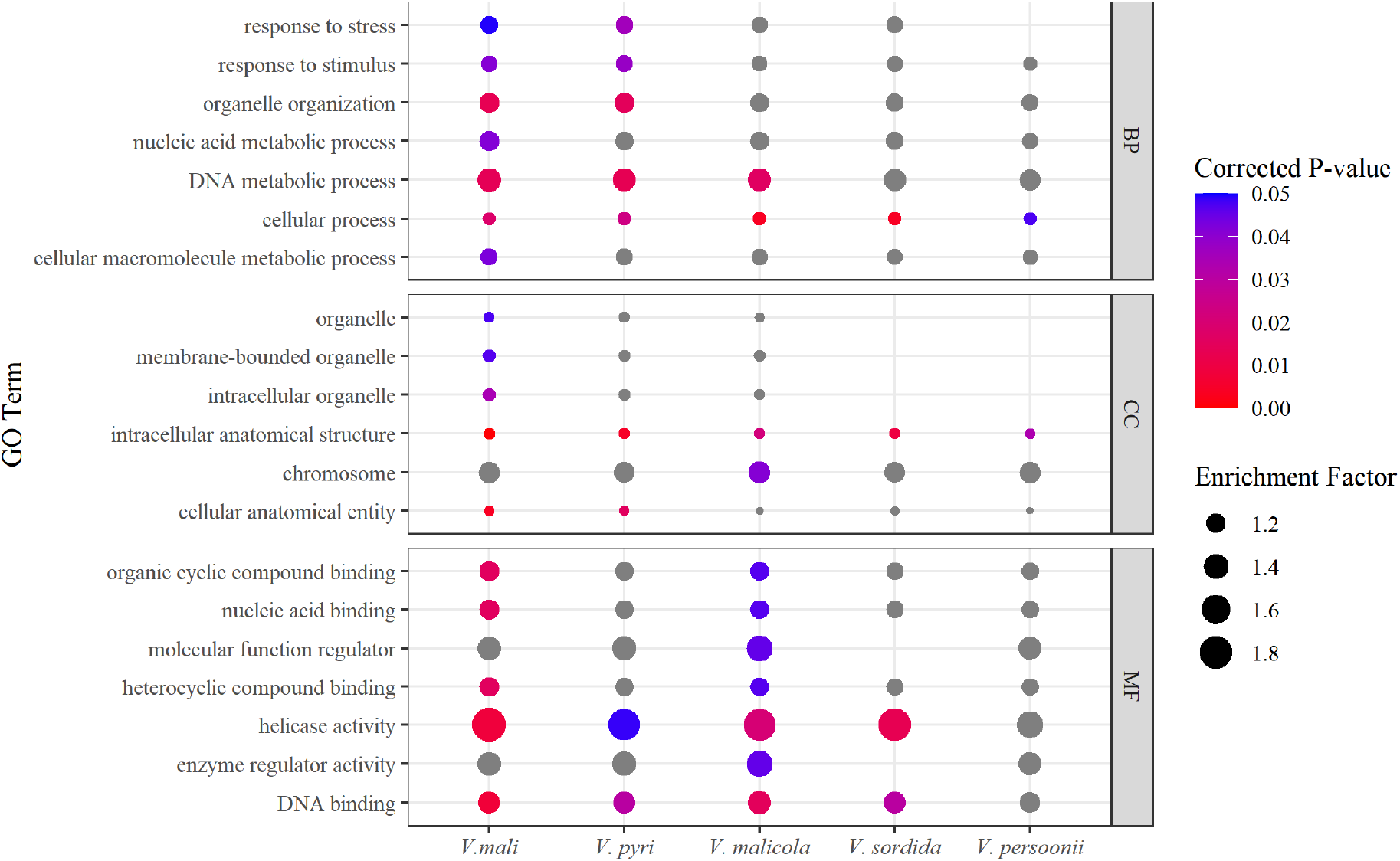
Gene ontologies over-represented in the genes under positive selection. (A) Gene ontology terms enriched, size indicate enrichment factor, corrected p values are color coded.

**Figure S7.**
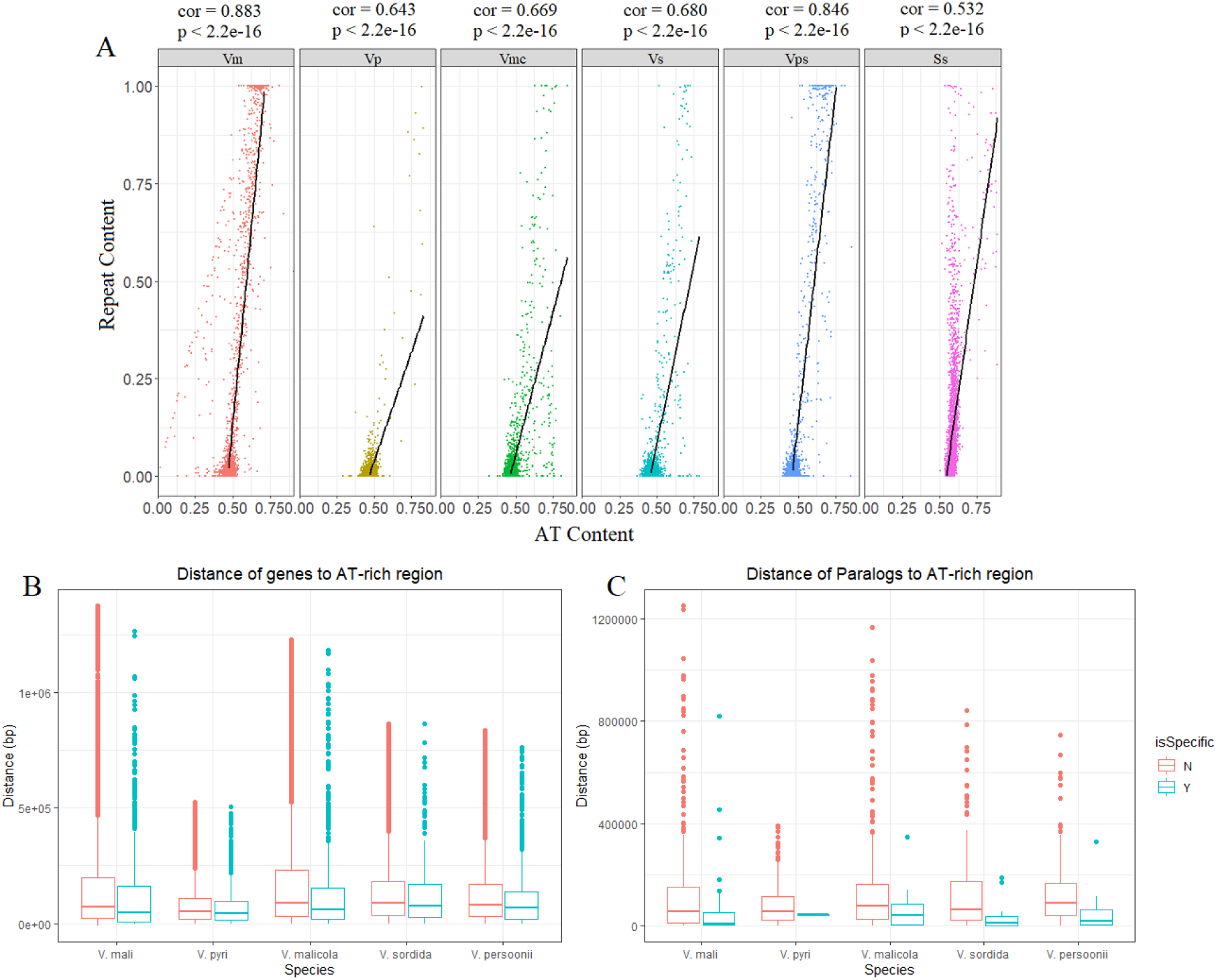
Correlation of repeat sequence and AT-rich region formation in valsa spp. (A) A significant coordination correlation between repeat elements and AT-rich regions was observed among all of the five Valsa species; (B) Positional correlation between AT-rich and species specific genes; (C) Positional correlation between AT-rich and species specific paralogs.

**Figure S8.**
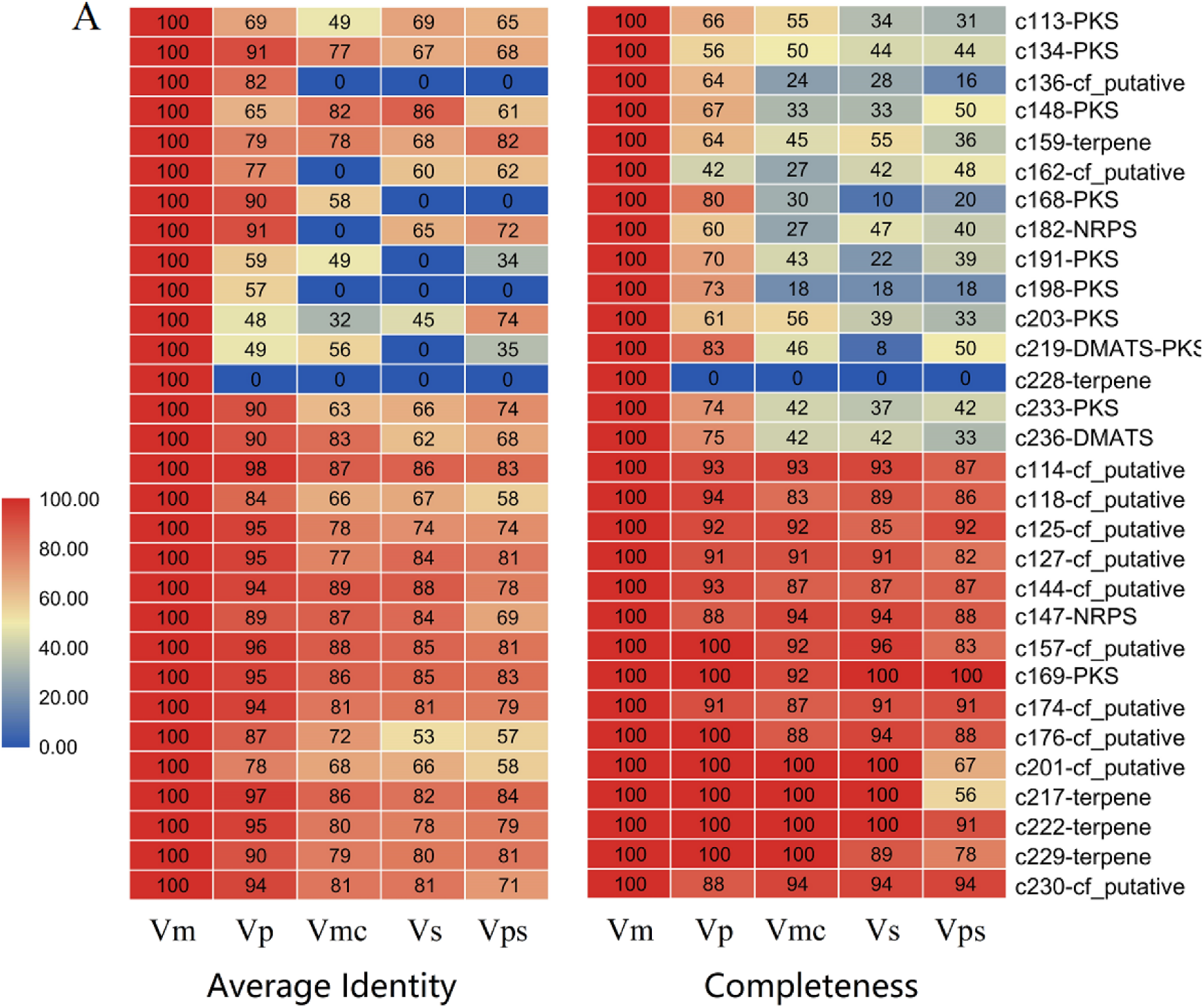
Conservation and diversification of secondary metabolic gene clusters among Valsa spp. (A) Average protein identity between SMs and their collinearity pairs in synteny blocks. (B) The ratio of the collinearity pairs and SM genes number.

## Supplementary Notes attached to this submission as separate files

- **Supplementary Data 1:** Summary of repeat contents of *Valsa Spp*..

